# MR-SP²: A Microreactor for Upward Pressure-Catapulting Laser Microdissection for Mass Spectrometry-Based Spatial Proteomics at Single-Cell Resolution

**DOI:** 10.64898/2025.12.23.695936

**Authors:** Manuel Metzger, Maximilian Maldacker, Tobias Hutzenlaub, Nils Paust, Oliver Schilling, Jan-Niklas Klatt

## Abstract

Laser capture microdissection (LCM) combined with liquid chromatography-tandem mass-spectrometry (LC–MS/MS) enables spatially resolved proteomics at few-cell scale, yet losses from minute LCM-cut specimens, particularly with upward pressure-catapulting systems and subsequent processing, limit depth and reproducibility.

We present MR-SP² (Microreactor-based Sample Preparation for Spatial Proteomics), a one-pot-in-solution workflow that integrates reproducible LCM-cut specimen capture, processing with minimized adsorptive losses, and pipetting-free transfer with Evotip disposable precolumns.

The workflow is demonstrated on a formalin-fixed paraffin-embedded (FFPE) murine kidney using upward pressure-catapulting LCM for targeted isolation of defined tissue specimens.

Across 50,000 µm³ regions (22 cells), MR-SP² modestly improved proteome coverage (3,381 ± 80 versus 3,174 ± 59 proteins). Decreasing sample input further accentuated the advantage of MR-SP² in maintaining higher identification rates, highlighting the successful reduction of adsorptive losses of the MR-SP² workflow. At 12,500 µm³ (5-6 cells), identifications increased to 1,145 ± 188 versus 302 ± 126. At 3,125 µm³ (1-2 cells), identifications reached 695 ± 112 versus 206 ± 51.

MR-SP² improves identification depth for few-cell FFPE samples nearly threefold and provides an LCM-compatible preparation that expands the robustness, applicability of upward pressure-catapulting LCM within the spatial proteomics toolkit.

## INTRODUCTION

Advances in highly sensitive, loco-regional liquid chromatography – tandem mass spectrometry (LC-MS/MS) proteomics have propelled the study of proteome biology in healthy and diseased tissues.^1,2^ By representing the tissue architecture, spatial proteomics enables precise mapping of protein distribution and abundance, thereby maintaining the spatial context critical for understanding tissue heterogeneity.^3–5^ Currently, mass spectrometry-based methods can identify hundreds to thousands of proteins from increasingly miniaturized samples including single cells.6 This capability provides a powerful phenotypic lens to dissect cellular heterogeneity, characterize rare populations, and discover spatially restricted biomarkers, offering fundamental insights for understanding disease mechanisms and developing targeted therapies.^7–9^

To translate these capabilities into practice, spatial proteomics workflows typically combine highly sensitive LC-MS/MS platforms with spatially resolved tissue sampling. In this context, laser capture microdissection (LCM) is the most widely used method for the excision-based, miniaturized dissection of tissue specimens for subsequent proteomic analysis. LCM uses a focused laser in combination with a microscope to excise and collect specific microregions from tissue with micrometer precision. Two LCM concepts are prevalent, featuring either gravity-based capture (dropping cut pieces downward) or upward pressure-catapulting.^10^ Those spatially defined LCM-cut specimens are then processed for mass-spectrometry-based proteomics. In practice, the achievable proteomic depth from such samples is predominantly limited by the sensitivity of the sample preparation which consists of LCM-cut specimen capture, sample processing (protein extraction, reductive alkylation, digestion, and acidification), and transfer to LC-MS/MS.

Losses during sample preparation from very small samples may result, amongst others, from:

i. partial or full loss of the LCM-cut specimen during transfer into and within the reaction vessel, especially for upwards-pressure catapulting LCM^10,11^;
ii. adsorptive losses of proteins and peptides to the inner surface of reaction vessels during extraction, reductive alkylation, and digestion^12^;
iii. pipetting steps for sample transfer to LC-MS/MS that increase adsorptive losses and lead to a substantial impact on proteomic depth^13^.

Overcoming these limitations is critical to fully harness the potential of spatial proteomics for widespread research and clinical applications.^14–16^ Existing approaches focused on minimizing adsorptive losses by employing droplet-based techniques^17,18^, hence avoiding excessive surface exposure of the sample^19^. An inherent problem of these approaches is increased evaporation, which creates variance in digestion.^20^ As evaporation is temperature dependent, temperature limits are imposed on open systems^3,18^ impairing protein extraction from FFPE samples^21^. Even though these approaches enable sample preparation with minimal adsorptive losses, reproducible handling of the nanoliter volumes requires complex dispensing systems^3,17^. As a result, conventional tube-based workflows remain widely used despite using larger volumes and multiple transfer steps, such as LCM-cut specimen transport within reaction vessels and pipetting into LC-MS/MS, which increase exposed surface area and consequently elevate adsorptive losses.^22,23^

For upward pressure-catapulting LCM, there is currently no workflow that addresses the discussed limitations in sample preparation. Hence, we developed the microreactor-based sample preparation for spatial proteomics (MR-SP²) workflow. MR-SP² centers on a novel microreactor that is compatible with upward pressure-catapulting LCM for reproducible capture of LCM-cut specimens. Sample processing within the microreactor is conducted in a closed, one-pot system, effectively minimizing evaporation and reducing adsorptive losses. Sample transfer to LC-MS/MS is performed by clipping the microreactor on an Evotip, hence avoiding pipetting of the sample and eliminating the associated adsorptive losses. Apart from the LCM and LC-MS/MS instruments, the workflow relies only on standard laboratory equipment and provides a replicable low-input spatial proteomics workflow.

## EXPERIMENTAL SECTION

### Animals and Ethics

Murine kidney tissue was obtained from C57BL/6 mice originally bred for the maintenance of the colony. The animals were housed under specific pathogen-free (SPF) conditions in accordance with the Federation of Laboratory Animal Science Associations (FELASA) guidelines^24,25^. Sacrifice and organ retrieval were performed for scientific purposes in strict compliance with the German Animal Welfare Act (Tierschutzgesetz §4 Abs. 3). The procedure was notified to and approved by the local Animal Welfare Officer (Tierschutzbeauftragter) and the responsible authorities (Regierungspräsidium Freiburg).

### Initial Tissue Processing

Following dissection, the kidney was fixed in 4% formalin (Sigma-Aldrich, St. Louis, USA) overnight. After fixation, the tissue was dehydrated through a graded ethanol series, cleared in a series of xylene (Carl Roth, Karlsruhe, Germany) washing steps, and infiltrated with molten paraffin wax. The paraffin-impregnated tissue was embedded in a paraffin block and stored at room temperature until further use. The tissue was then cut in 5 µm thick sections and mounted on PEN slides (Carl Zeiss Microscopy, Jena, Germany) for laser microdissection. Before conducting LCM, the section was deparaffinized in consecutive steps of 99%, 96% 70% xylol and 50% ethanol (Carl Roth) and a final wash procedure in 70% ethanol (Carl Roth) before being rehydrated and kept in water until dissection.

### Manufacturing of the Microreactor

The microreactor (Figure 1A) was manufactured using thermoforming.^26,27^ For this, the design was created using SolidWorks 2021 (Dassault Systèmes, Vélizy-Villacoublay, France) and subsequently milled in polymethyl methacrylate (PMMA) using a Kern Micro HD (Kern Mikrotechnik, Hohenlohe, Germany) milling machine. The PMMA master was molded using Elastosil RT607 polydimethylsiloxane (PDMS, Wacker Chemie, München, Germany). Thermoforming was performed on a hot embossing machine (Wickert Maschinenbau, Landau in der Pfalz, Germany) using the PDMS mold and 2x 400 µm COC foil (TekniPlex, Wayne, USA) consisting of a low-melting side (COC 8007) and a high-melting side (COC 6013). After thermoforming, the microreactor is cut out using a laser cutter (Universal Laser Systems, Scottsdale, USA).

**Figure 1.**
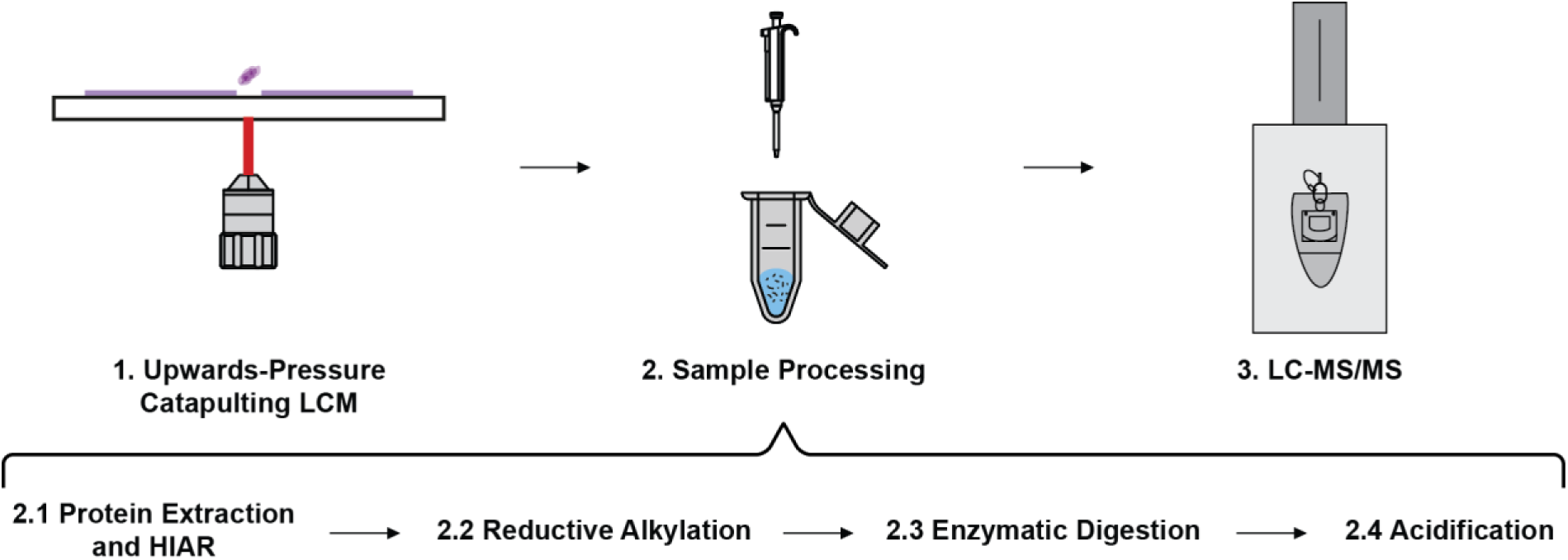
The schematic process of sample preparation for spatial proteomics deploying upwards-pressure catapulting LCM: (1) LCM; (2) sample processing which is further split into protein extraction and heat-induced antigen retrieval, reductive alkylation, enzymatic digestion, and acidification; (3) LC-MS/MS analysis.

For manufacturing of the microreactor cap (Figure 1B), a 3D printing process using the Arburg Freeformer 300-3X (Arburg, Lossburg, Germany) was employed using APEL COC 6013T (Mitsui Chemicals, Minato, Japan) as a material and Armat 11 (Arburg) served as support material during the printing process. A cylinder with a diameter of 15 mm and a thickness of 4 mm was printed. After the print, a hole with a depth of 2 mm was drilled into the cylinder using a 5.6 mm drill bit. To prevent thermo-bonding of the microreactor and the cap of the microreactor during the 95 °C extraction step, the drill hole was coated with a 10% w/w solution of PTFE (DuPont Fluoroproducts, Wilmington, USA) in Fluorinert FC-40^28^ (Sigma-Aldrich) and the Fluorinert was subsequently evaporated. The holder for the microreactor (Figure 1B) was also 3D printed using the same printing process as for the cap.

### Laser Capture Microdissection

Laser capture microdissection was conducted using the Zeiss PALM Microbeam platform connected with a Zeiss Microscope (MBC01092), Control Box (2_Rev. 4.2), a Stage (Stage IIB) and the capture device (RoboMover) mounted with the Collector Set SingleTube II 500. The specimens were cut with CryLaS UV-Laser and controlled with the software installations of PALMRobo (V 4.6.0.4), MTB (V 1.8.1.5), AxioVision (4.8.3.0) and ControlBox (1,170,146M).

The tissue was dried before installing the tube or microreactor. To prevent biological intra-tissue variance overshadowing the technical variance, we selected homogenous renal corticoid tissue sections (Supplemental Figure 1, A+B). The tissue was dissected in “Cut” mode for the given tissue area with different sizes before transferring the LCM-cut specimen to the reaction vial with a series of LPC shots (Supplemental Figure 1, C-E).

For conventional tube capture, 200 μL PCR tubes were used (PCR microcentrifuge tube PP, 0.2 mL, 8-strips, Nerbe Plus, Winsen, Germany). For capture, the caps of the tubes were filled with 40 μL acetonitrile. The successful transfer in the acetonitrile volume was confirmed visually before starting the sample processing. A subsequent transfer step of the acetonitrile and LCM-cut specimen from tube cap to tube body was performed by short centrifugation using a mini centrifuge with a maximum of 2000 g.

For microreactor capture, due to lower cavity volume the microreactor was filled with 20 μL acetonitrile. The microreactor was placed in the position taken by the cap for conventional tube capture.

In both cases, the acetonitrile was evaporated by centrifugal vacuum evaporation after capture using a Speedvac (Eppendorf, Hamburg, Germany).

### Sample Processing for LC-MS/MS

An adapted version of the one-pot workflow for few-cell proteomics by Tsai et al.^29^ was performed. 5.0 µL of 0.2 % Dodecyl-β-D-maltosid (DDM, Carl Roth) in 5.0 mM Tris(2-carboxyethyl)phosphine hydrochloride (TCEP, Sigma-Aldrich) were added with incubation at 95 °C for one hour followed by addition of 1.0 µL of 60.0 mM chloracetamide (CAA, Sigma Aldrich) in 5.0 mM TCEP (final CAA concentration of 10 mM) at 37 °C for 30 minutes. Then, 1.0 µL of 175 mM ammonium bicarbonate (Sigma-Aldrich) in 5 mM TCEP were added followed by 1.0 µL of a trypsin (50 ng/µL, Serva) and LysC (25 ng/µL, Serva, Heidelberg, Germany) solution. Enzymatic digestion was carried out for 16 hours at 37 °C. Samples were then acidified by addition of 12 µL of 10 % formic acid (FA, Sigma-Aldrich).

### LC-MS/MS

Evotips were preconditioned according to manufacturer guidelines. Samples were transferred to Evotips (Evosep, Odense, Denmark) using either manual pipetting (for conventional tube capture) or by centrifugal transfer (microreactor clipped on Evotip, see Supplemental Figure 4) and centrifuged for one minute at 800 g. The samples were loaded with a volume of 20 µl and washed twice with 20 µl 0.1% FA in H2O. The samples were kept in 0.1% FA in H2O at 4 °C until MS-acquisition. Peptides were separated on a EvoSep Performance 15 µm column (EV1137) tempered to 40 °C with a 30 SPD gradient and injected utilizing the captive spray source with a 20 µm ZDV sprayer (Bruker Deltatonics, Billerica, USA).

MS/MS acquisition was conducted in diaPASEF mode with an accumulation and ramp time of 150 ms. Precursors within the ion mobility range of 0.6-1.6 1/K0 and within the m/z range of 400-1200 Th were selected for fragmentation with one MS1 cycle followed by 32 MS2 scans from two PASEF cycles each. Precursors were fragmented with a fragmentation energy of 20-59 eV increasing linearly with ion mobility and MS2 spectra were acquired in a range of 100-1700 Th.

### Data Analysis

LC-MS/MS data was searched with DIA-NN 2.2^30,31^ against a murine proteome database (2023, 20591 entries) appending iRT peptides reannotated and modified by the addition of common contaminants. A spectral library was predicted from the given fasta file utilizing tryptic peptides and allowing for a maximum of 1 missed cleavage. Precursors with a peptide length between 7-50 Th within a MS1 range of 400-1200 Th were matched with fragment ions within a MS2 range of 100-1700 Th. N-terminal M excision and oxidation at M were set to variable modifications, carbamidomethylation at C was set as fixed modification. Protein inference was activated, scoring was run in generic mode, NNs were selected for learning with QuantUMS32 and conducted in high precision mode. Only identifications with a q-value < 0.01 were retained for analysis. Identification numbers were reported from two separate DIA-NN analyses without and with applying MBR, with MBR being confined to LC-MS/MS analyses of equal input amounts. Protein abundances were determined using DIANN’s integrated MaxLFQ normalization, and peptide quantification was based on the Precursor.Quantity metric. Data analysis was done using an in-house python script including packages shown in the Supporting Information (see Data Availability). Statistical significance was evaluated with a Mann-Whitney U test for all pairwise comparisons except for the comparison of average peptide lengths where a Student’s t-test was used.

## RESULTS AND DISCUSSION

### The MR-SP² Workflow Mitigates Sample Loss in Upwards-Pressure Catapulting LCM

The microreactor (Figure 2A+B) is the core component of the MR-SP² workflow (Figure 2C), designed to address the limitations in LCM-cut specimen capture, sample processing, and transfer to LC-MS/MS.

**Figure 2.**
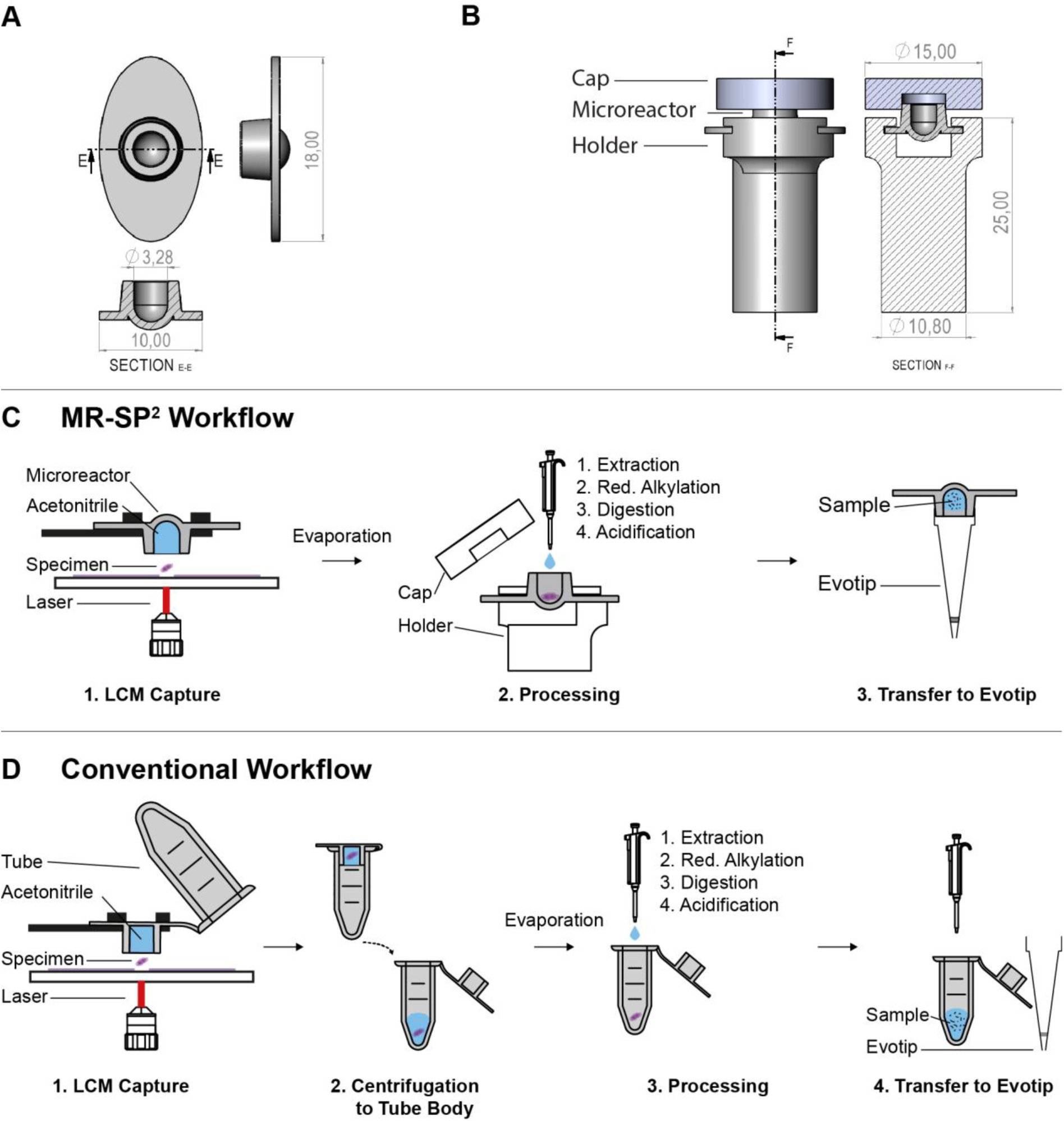
**(A)** Technical drawing of the microreactor; all dimensions in mm. **(B)** All three components assembled together with the microreactor in the holder and the cap attached to avoid evaporation; all dimensions in mm. **(C)** MR-SP² workflow: (1) The microreactor filled with 20 µL acetonitrile captures an upward pressure-catapulted LCM-cut specimen, is subsequently mounted in a 3D-printed holder, and the acetonitrile is evaporated in a SpeedVac under rotation. (2) After evaporation, a closed one-pot in-solution workflow (extraction, reductive alkylation, digestion, acidification) is performed, with a polymer cap applied after each step to prevent evaporation. (3) The microreactor is clipped onto an Evotip, enabling pipetting-free transfer to LC–MS/MS. **(D)** Reference tube-based workflow: (1) A tube cap filled with 40 µL acetonitrile captures an LCM-cut specimen by upward pressure-catapulting. (2) Specimen and acetonitrile are centrifuged to the tube body and acetonitrile is evaporated in a SpeedVac under rotation. (3) After evaporation, a closed one-pot workflow (extraction, alkylation, digestion, acidification) is performed, with the tube cap applied after each step to prevent evaporation. (4) The sample is pipetted into an Evotip for LC-MS/MS analysis.

The microreactor is fully compatible with the Zeiss PALM Microbeam platform and its RoboMover for upward-pressure catapulting LCM. To avoid specimen loss due to adherence to capture adhesives^33^, fluid-assisted collection^17^ with acetonitrile is employed. The microreactor requires only 20 µL acetonitrile for specimen capture, simplifying localization and visual verification compared to the higher volume of the conventional workflow (40 µL capture volume, Figure 2D).

By capturing LCM-cut specimens directly in the microreactor, post-dissection tissue transport is eliminated for the MR-SP² workflow. Conventional workflows, however, require an additional centrifugation step to transfer specimens from the tube cap to the tube body, which resulted in specimen loss even after confirmed initial capture (Supplemental Figure 2, 22 cells – Conventional – R6). To conclude the capture, the specimen is localized at the base of the reaction vessel by solvent evaporation under rotation.

For sample processing, the microreactor’s cavity volume is optimized to a total workflow volume of 20 µL, minimizing the area exposed to the inner surface of the reaction vessel. It can withstand temperatures above 95 °C, enabling high-temperature protein extraction of FFPE tissues. To prevent evaporation and preserve sample integrity, the microreactor is sealed with a cap (Figure 2B) before the incubation steps.

To enable pipetting-free sample transfer to LC-MS/MS, the microreactor is designed to clip onto an Evotip, significantly reducing adsorptive losses associated with pipetting steps. Both the MR-SP² and conventional workflows use a total processing volume of 20 µL (see Experimental Section). By minimizing the exposed surface area and eliminating pipetting during sample transfer, MR-SP² reduces the cumulative surface exposure by 64% compared to the conventional workflow (33.4 mm² vs. 92.8 mm², Supplemental Figure 3), thereby substantially decreasing non-specific protein and peptide binding.

### MR-SP² Yields Improved Proteome Coverage and Delivers High-Quality Data

We evaluated capture efficiency for conventional tubes and the microreactor by LCM of 100×100 µm² regions from 5 µm murine kidney sections in 16 replicates each. Visual inspection confirmed successful specimen capture in 15 of 16 microreactor-captured samples (94%), whereas only 12 of 16 samples (75%) were successfully captured using the conventional tube-based workflow. In addition, the two-fold reduction in capture volume in the microreactor facilitated visual confirmation of specimen presence, enabling more reliable verification compared to the tube-based workflow.

LCM-cut specimens were further processed and analyzed by LC-MS/MS to examine whether the MR-SP² workflow approach translates into deeper proteomic coverage (Figure 2B+C). We are reporting identification numbers without and with applying MBR, with MBR being confined to LC-MS/MS analyses of equal input amounts. Notably, one conventionally processed sample yielded markedly low identifications (515 peptides and 219 proteins) despite confirmed dissection and transfer into the tube cap (Supplemental Figure 2). By facilitating visual confirmation of specimen presence and eliminating subsequent transfer steps within the microreactor, the MR-SP² workflow features an increased overall procedural robustness. In order to evaluate the workflow performance, the low identification sample of the conventional workflow was not included in the further analysis.

MR-SP² demonstrated superior sensitivity across different input amounts (Figure 3A+B). For a dissected volume of 50,000 µm³ (22 cells^34,35^), it yielded an average of 16,021 ± 1,009 peptide identifications (20,229 ± 895 with MBR; n = 11), a significant increase over the 13,514 ± 740 (16,852 ± 513 with MBR; n = 7) identified peptides from the conventional workflow (Figure 3A; p < 0.01, Mann-Whitney U test). The improved sensitivity was less pronounced on the protein level with an increase of approximately 6% more proteins (Figure 3B; p = 0.051, Mann-Whitney U test); indicative of improved sequence coverage of individual proteins. We further probed reduced input amounts in addition to the 50,000 µm³ LCM-cut specimens: 12,500 µm³ (5-6 cells^34,35^) and 3,125 µm³ (1-2 cells^34,35^). For an input of 5-6 tissue cells, the MR-SP² (n=5) yielded an average of 3,467 ± 902 peptide identifications (4,495 ± 912 with MBR); corresponding to a > 5-fold increase as compared to conventional workflow (n = 5), which yielded 685 ± 420 peptide identifications (801 ± 380 with MBR) (p < 0.05, Mann-Whitney U test). Unlike for the 22-cell cell input, the improved peptide coverage also translates to a strongly improved protein coverage: the MR-SP² workflow resulted in an average of 914 ± 216 (1,145 ± 188 with MBR) protein identifications, which is > 3-fold more than the 231 ± 135 (302 ± 126 with MBR) proteins identified by the conventional workflow (p < 0.05, Mann-Whitney U test). For an input of 1-2 tissue cells, the microreactor workflow (n = 5) yielded an average of 1,841 ± 480 peptide identification (2,320 ± 423 with MBR); corresponding to a > 3-fold increase as compared to conventional workflow (n = 5), which yielded 541 ± 170 peptide identification (601 ± 111 with MBR) (p < 0.05, Mann-Whitney U test). The improved peptide coverage also translates to improved protein coverage: the MR-SP² workflow resulted in an average of 557 ± 135 (695 ± 112 with MBR) protein identifications, which is > 3-fold more than the 176 ± 73 (206 ± 51 with MBR) proteins identified by the conventional workflow (p < 0.05, Mann-Whitney U test).

**Figure 3.**
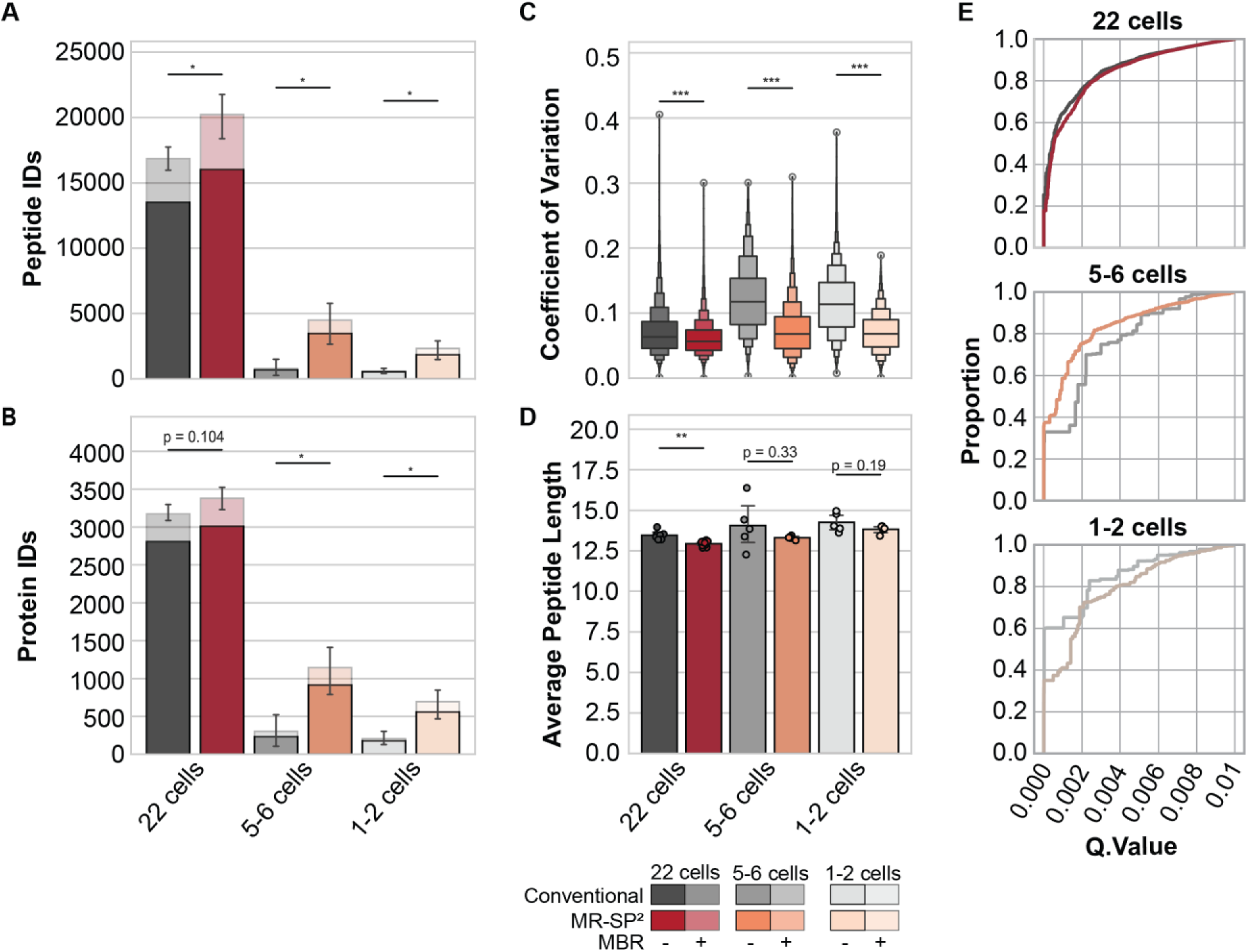
**(A)** Number of identified peptides and **(B)** proteins from 22, 5-6, and 1-2 cells of a FFPE murine kidney slide. Saturated bars denote identifications without applying the match between runs algorithm (MBR), while lighter bars indicate additional matches assigned by MBR within the same condition. Error bars represent the standard error of the mean for n=7, n=11 and n=5 samples for 22 cells - conventional, 22 cells - MR-SP² and the remaining conditions, respectively. Significance was tested using a Mann-Whitney U test on identifications with MBR (*: p < 0.05, **: p < 0.01). **(C)** Coefficient of variation for the log₂ transformed precursor intensity among peptides identified in more than 3 samples per condition (**: p < 0.01, ***: p < 0.001, Mann-Whitney U test). **(D)** Peptide length in amino acids. Significance was tested with Student’s t-test, ** depicts p < 0.01. **(E)** Q.Value distribution for different input amounts in cell equivalents for MR-SP² and conventional sample preparation.

Previous observations already reported that an increased surface area causes significant sample losses due to adsorption with decreasing sample inputs being affected disproportionately^36^. In line with this, we observed a significant increase in peptide and protein identifications, especially for reduced input levels, and link this to the omission of one pipetting step in MR-SP².

To determine the coefficient of variation (CV) of the precursor intensities per processing, we selected precursors with at least three valid values per condition. At 22 cells (sampled from comparable kidney areas), both workflows indicated a high quantitative accuracy with mean CVs of 0.07 ± 0.036 from 21,018 and 0.061 ± 0.027 from 29,246 precursors for the conventional and MR-SP² workflow, respectively. In addition, the majority of all precursors yielded a CV < 0.15 (Figure 3C). A multi-laboratory assessment of reproducibility, qualitative, and quantitative performance of LC-MS/MS based proteomics suggested bioanalytical acceptance of < 20% CV with our CV values comparing favorably^37^. Quantitative accuracy remained stable even under extremely low input. At 5–6 cells, variability fell from 0.12 ± 0.06 based on 462 precursors to 0.07 ± 0.04 based on 5,171 precursors. At 1–2 cells, it decreased from 0.116 ± 0.051 on 566 precursors to 0.071 ± 0.031 on 2,774 precursors (Figure 3C, p < 0.001). This substantial reduction in variance highlights the robustness of MR-SP² and its suitability for reproducible analysis of near single-cell proteomes. The data suggest that by the elimination of one transfer step and the reduction of the exposed total surface entropic peptide-level losses can be omitted that would otherwise affect reproducible quantification across several LC-MS/MS runs.

In addition, we determined the average length of the identified peptides per LC-MS/MS run (Figure 3D). For the input amount of 22 cells, MR-SP² derived peptides showed a minor (less than one residue on average), albeit significant tendency for reduced amino acid length (12.9 ± 0.05 versus 13.5 ± 0.09; p < 0.01, Student’s t-test). A similar, yet not significant trend was observed for the reduced input amount of 5-6 cells with 13.3 ± 0.05 and 14.1 ± 0.7 amino acids in length for MR-SP² and the conventional workflow, respectively (p = 0.33, Student’s t-test). Similarly, LC-MS/MS runs from 1-2 cells exhibited a similar average peptide length with 13.8 ± 0.1 and 14.3 ± 0.3 amino acids in length. Overall, MR-SP² peptides show only a marginal reduction in average length compared to the conventional workflow. Importantly, this minor difference does not produce a consistent shift across LC-MS/MS runs, indicating that peptide size distributions are largely maintained even at minimal input levels.

Furthermore, we employed the Q.Value metrics of DIA-NN as a parameter for spectral quality of sequence identifications (Figure 3E). No major differences between the conventional and MR-SP² workflow are evident. This suggests that the increased identifications observed in the MR-SP² workflow are not driven by lowered confidence thresholds or increased false positives but rather reflect true improvements in recovery and sensitivity.

### Gains in MR-SP² Originate from Low-Abundant Peptides

We assessed to which extent each workflow is contributing to the identification of specific peptide subsets. To this end, we used the accumulated peptide identifications per workflow, including peptides identified in a single LC-MS/MS run with a global q-value < 0.01. For all three input amounts, there was a sizable proportion of peptides uniquely identified in either the conventional or the MR-SP² workflow (Figure 4).

**Figure 4.**
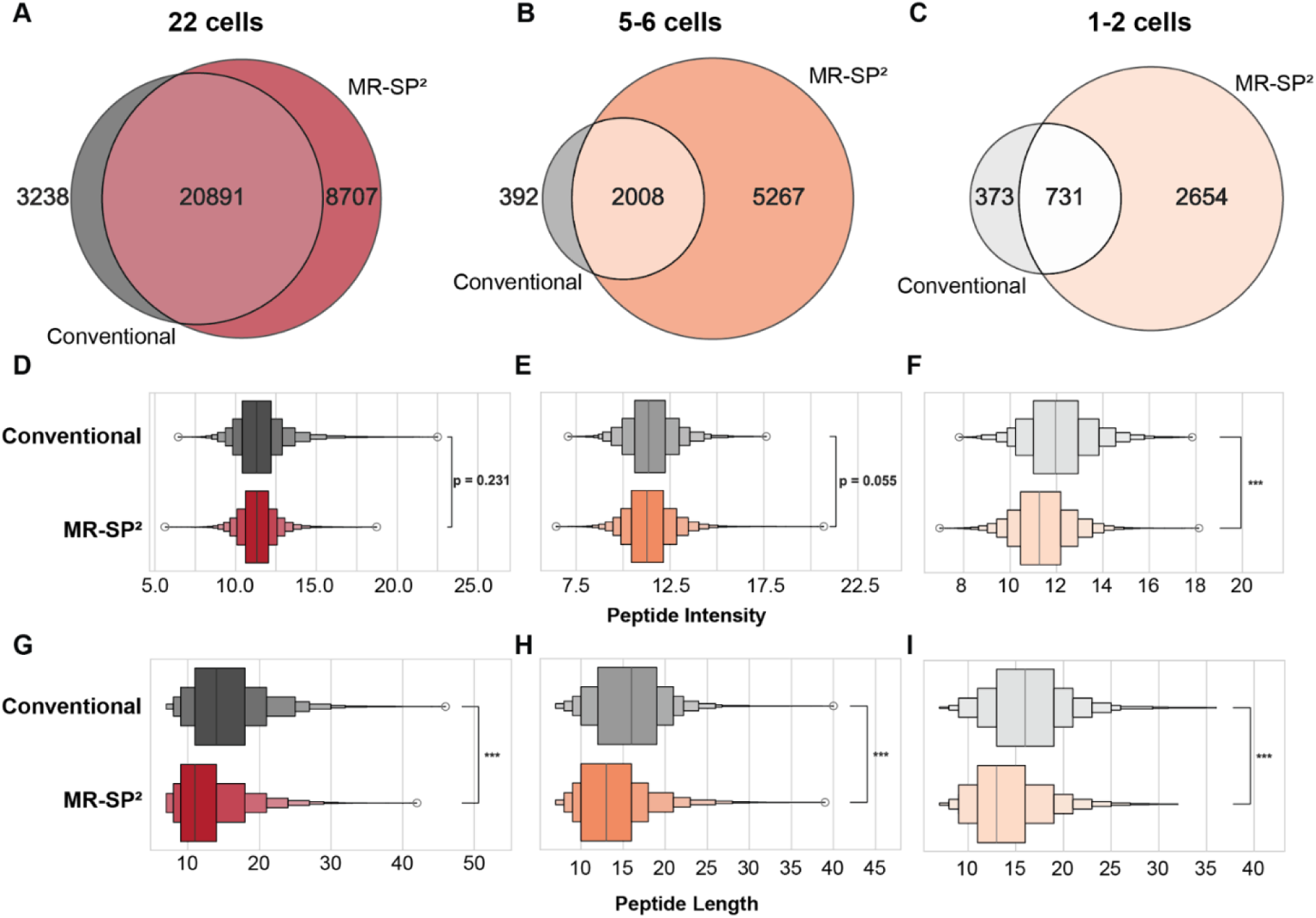
Cumulative Venn diagram of all identified peptides for MR-SP²-based sample preparation or conventional sample preparation resulting from 22 **(A)**, 5-6 **(B)** or 1-2 **(C)** cells from FFPE murine kidney being processed. MBR was activated across all samples from the same condition. Log2 peptide intensity of unique peptides from A-C for either the MR-SP²- or conventional workflow from 22 **(D)**, 5-6 **(E)** or 1-2 **(F)** cell equivalents being processed. Peptide length of unique peptides for either the MR-SP² or conventional workflow from 22 **(G)**, 5-6 **(H)** or 1-2 **(I)** cells being processed. Identification, peptide intensity and peptide length from n=7, n=11 and n=5 samples for 22 cells - Conventional, 22 cells - MR-SP² and the remaining conditions, respectively. Significance was tested with Mann-Whitney U test (ns: not significant, ***: p < 0.001).

For unique peptides from MR-SP²–processed samples, we obtained consistently lower MS2-level intensities compared to peptides unique to the conventional workflow, a trend that became most pronounced at the lowest input amounts (Figure 4, E-F). At 22 cells, the mean log₂ intensity was 11.3 ± 1.2 for MR-SP² and 11.4 ± 1.6 for the conventional workflow (*p* = 0.2, Mann-Whitney U). At 5–6 cells, intensities were 11.4 ± 1.3 versus 11.5 ± 1.4 (*p* = 0.055, Mann-Whitney U), and at the 1–2 cell level, 11.35 ± 1.34 versus 12.04 ± 1.56 (*p* < 0.01, Mann-Whitney U). Thus, the data showcase the conceptual reduction of exposed surface area in MR-SP² is enabling the detection of low abundant unique peptides that would otherwise be below the limit of detection and are prone to adsorptive losses^38,39^.

We also determined the observable peptide sequence length (Figure 4, G-I). At 22 cells, MR-SP² peptides were shorter on average (11.3 ± 1.2 amino acids) than those from the conventional workflow (14.8 ± 5.5; *p* < 0.001, Mann-Whitney U). This difference persisted at 5–6 cells (13.4 ± 4.4 vs. 15.7 ± 4.7; *p* < 0.001, Mann-Whitney U) and was also apparent at 1–2 cells (13.8 ± 4.2 vs. 16.0 ± 4.6; *p* < 0.001, Mann-Whitney U). The decrease in peptide length as an indicator for better digestion efficiency^40^ hereby demonstrates that the MR-SP² contributes to enhanced cross LC-MS/MS run quantification of low abundant proteins inferred from low abundant peptides.

Collectively, these results show that albeit not all comparisons reach statistical significance, differences in intensity and peptide length become more prominent with lower sample input. The consistent trend toward shorter and less intense peptides in the MR-SP² workflow indicates improved transfer of low abundant peptides from the sample to the instrument due to reduced absorption and transfer-induced sample loss.

## CONCLUSION

The MR-SP² workflow introduces a microreactor-based sample preparation approach that effectively addresses the principal bottlenecks - sample capture, adsorptive losses, and inefficient extraction - in spatial proteomics sample preparation, particularly for upward pressure-catapulting LCM platforms (e.g., Zeiss PALM Microbeam).

By enabling transfer-free, pipetting-free, one-pot in-solution sample processing, MR-SP² significantly reduces specimen loss and adsorptive losses, resulting in improved sensitivity and reproducibility for low-input and near single-cell proteomics. Demonstrated gains in protein identification and quantitative robustness across a range of tissue inputs highlight its suitability for deep characterization of spatially defined tissue regions.

The replicability of the design and compatibility with standard laboratory equipment further enhances accessibility, supporting broad adoption in spatial proteomics research and clinical applications. MR-SP² expands the spatial proteomics toolkit and facilitates more comprehensive understanding of tissue biology at cellular resolution.

### Declaration of Generative AI and AI-assisted Technologies in the Writing Process

Manuscript editing was supported by Gemini (2.5 Flash, Google) and ChatGPT (OpenAI). The authors reviewed and edited the content and take full responsibility for the content of the published article. Development of data analysis scripts (python) was supported by ChatGPT (OpenAI).

### Data Availability

Raw mass spectrometry data have been deposited in MassIVE under accession **MSV000099955** *(available via MSV000099955_reviewerM; password: MR-SP2025_)*. Analysis scripts are accessible through a private GitHub link (temporary access provided for peer review):

https://github.com/MaldackerM/MR-SP-A-Microreactor-for-Laser-Microdissection-for-Single-Cell-Spatial-Proteomics.git.

## AUTHOR INFORMATION

### Corresponding Author

Correspondence to Jan-Niklas Klatt and Oliver Schilling.

### Author Contributions

The manuscript was written through contributions of all authors. All authors have given approval to the final version of the manuscript. †,* These authors contributed equally.

### Funding Sources

This work was supported by the State of Baden-Württemberg through the Invest BW innovation program (Grant name: KASPAR, project number BW1_1198/02 and BW1_1198/03). The funding agency had no role in study design, data acquisition, data interpretation, or manuscript preparation. The authors acknowledge additional institutional support from the Medical Center Freiburg and the Department of Microsystems (IMTEK), Freiburg.

O.S. acknowledges funding by the German Research Foundation (projects 446058856, 438496892, 524803248, 546330039, 431984000 (SFB 1453 “NephGen”), 441891347 (SFB 1479 “OncoEscape”), 423813989 (GRK 2606 “ProtPath”), 322977937 (GRK 2344 “MeInBio”), 507957722), the ERA PerMed program (PerCareGlio), the ERA TransCan program (BMBF 01KT2201 “PREDICO”, 01KT2333 “ICC-STRAT”), the German Consortium for Translational Cancer Research (project “Impro-Rec”), the investBW program (project BW1_1198/03 “KASPAR”), the BMBF KMUi program (project 13GW0603E, project “ESTHER”), and nanodiag (03ZU1208AA nanodiag BW).

## ACKNOWLEDGMENT

Technical contributions from the Prototyping Workshop at Hahn-Schickard in Microreactor manufacturing are gratefully acknowledged. This work received financial support from the State of Baden-Württemberg (Invest BW, Project KASPAR), the DFG, and the DKTK/DKFZ. We further thank Carl Zeiss Microscopy GmbH for their collaboration and support of the LCM platform.

## ABBREVIATIONS

CAA,: chloroacetamide;
COC,: cyclic olefin copolymer;
CV,: coefficient of variation;
DDM,: odecyl-β-D-maltoside;
FA,: formic acid;
FFPE,: formalin-fixed paraffin-embedded;
LC-MS/MS,: liquid chromatography–tandem mass spectrometry;
LCM,: laser capture microdissection;
MBR,: Match Between Runs;
MR-SP²,: Microreactor-based Sample Preparation for Spatial Proteomics;
PMMA,: polymethyl methacrylate;
PDMS,: polydimethylsiloxane;
TCEP,: Tris(2-carboxyethyl)phosphine hydrochloride.

## Notes

### Competing Interest Statement

The authors have declared no competing interest.

