## Supplemental for "MR-SP²: A Microreactor for Upward Pressure-Catapulting Laser Microdissection for Mass Spectrometry-Based Spatial Proteomics at Single-Cell Resolution"

ASSOCIATED CONTENT

### Supporting Information

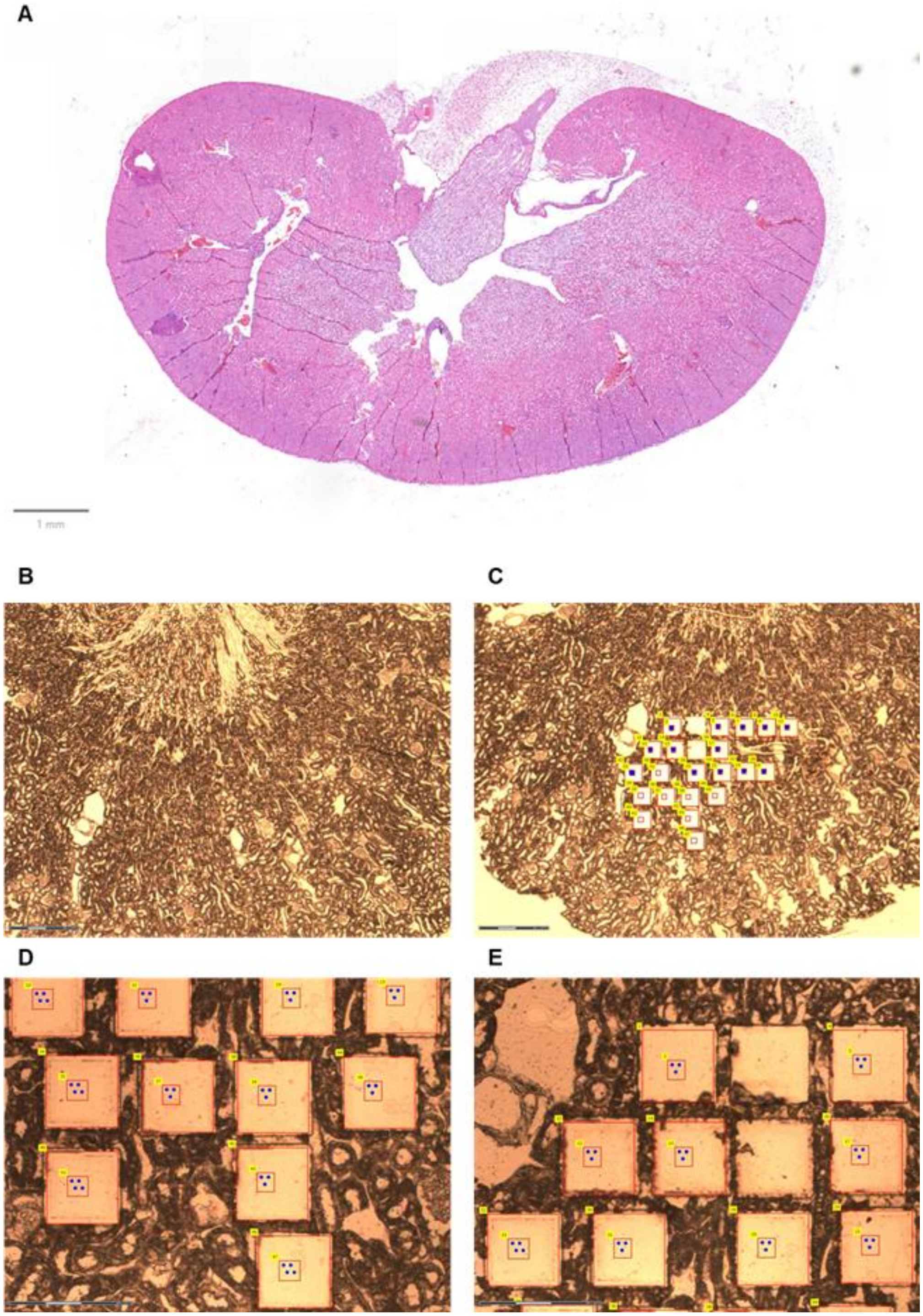

***Supplemental Figure 1.*** ***(A)*** *High resolution scan of the murine kidney section after hematoxylin-eosin staining. The scale bar represents 1mm. Microscopic image of the dissected region from deparaffinized murine kidney before* ***(B)*** *and after* ***(C)*** *laser microdissection. The scale bar represents 300 µm. Lasermicrodissected regions in 20-fold magnification* ***(D)*** *and* ***(E)*** *with the scale bar representing 150 µm.*

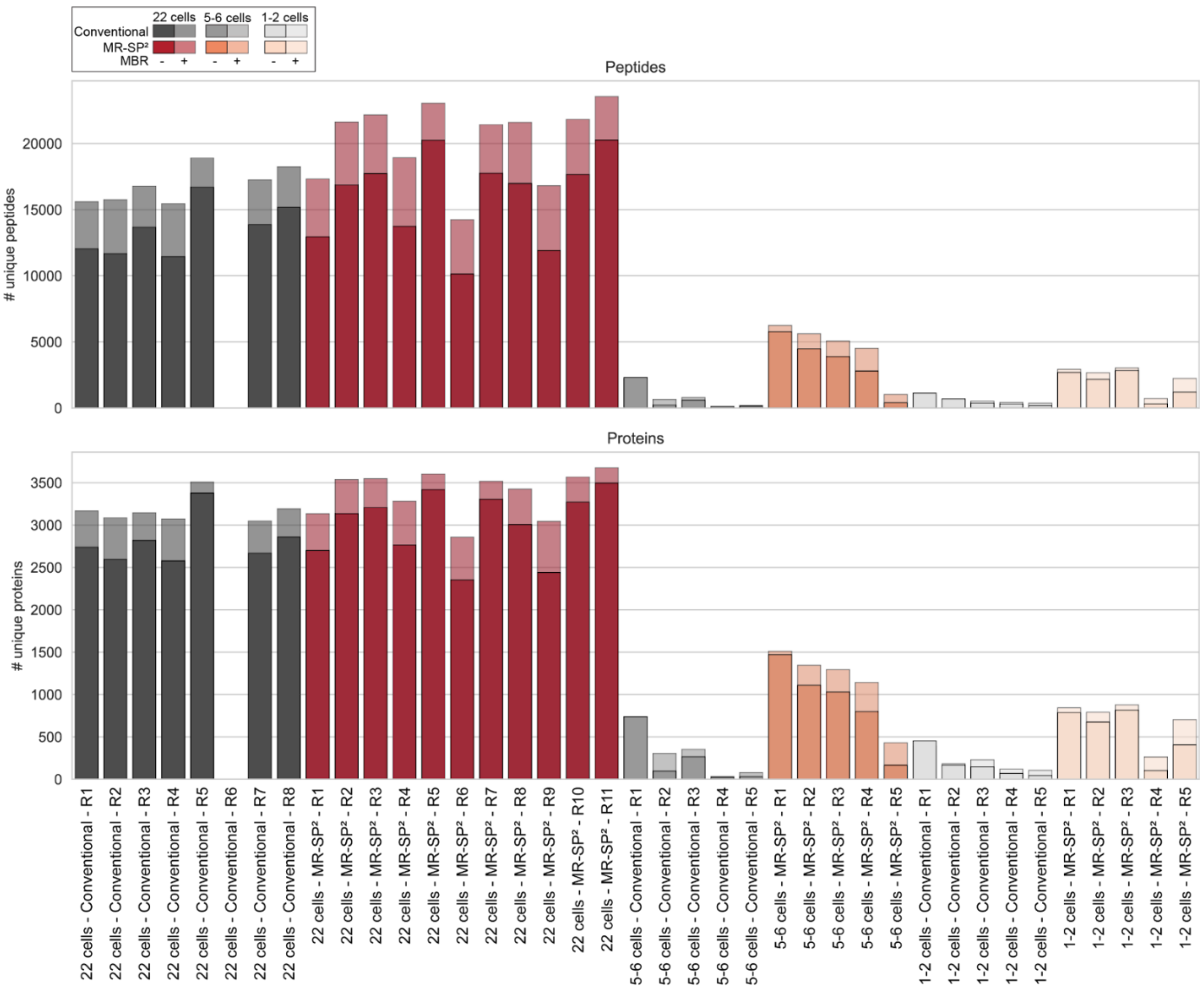


***Supplemental Figure 2.*** *Number of identified peptides and proteins from 22, 5-6 and 1-2 cells of a formaline-fixed murine kidney slide for individual samples. Saturated bars represent identifications without MBR activated, while lighter bars indicate additional identifications attributed to MBR across samples from the same condition.*

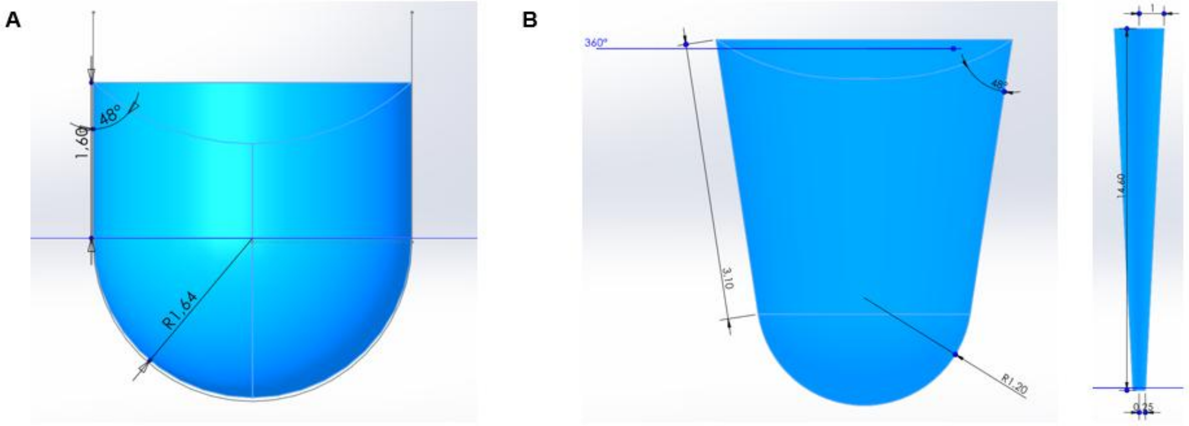


***Supplemental Figure 3. (A)*** *Depicted is the CAD model of the 20 µL sample volume after complete sample processing before transfer to the Evotip. The model shows the exposed polymer surface area to the sample that is responsible for adsorptive losses. The surface area is 33.4 mm². All measurements shown are in mm.* ***(B)*** *Depicted is the CAD model of the 20 µL sample volume after complete sample processing before the transfer to the Evotip (left) in a 200 µL microtube. Additionally, the model on the right shows the 20 µL sample volume in a 100 µL pipette tip that is necessary for transfer to the Evotip. The cumulated surface area of sample processing and pipetting transfer to the Evotip is 92.8 mm². All measurements shown are in mm.*

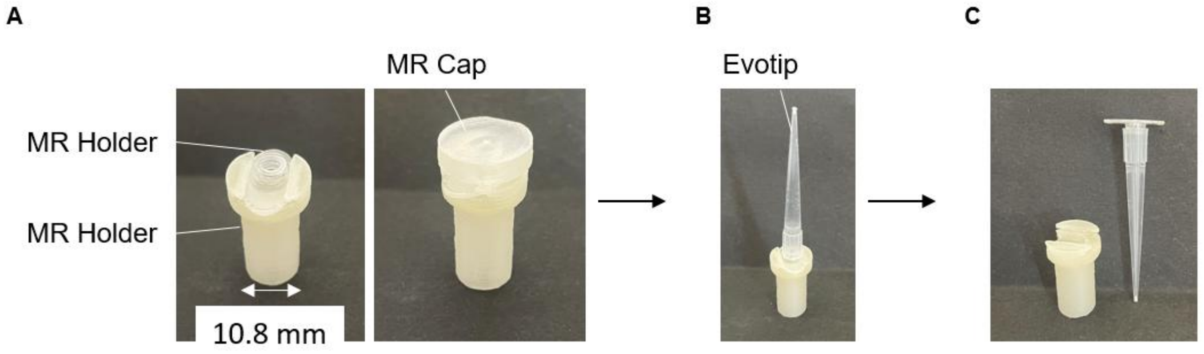


***Supplemental Figure 4. (A)*** *The picture on the left side shows the microreactor placed in the holder for sample processing. The picture on the right side shows the applied microreactor cap*

*when the microreactor is closed.* ***(B)*** *The picture shows the microreactor still in the holder and an Evotip is clipped onto the microreactor.* ***(C)*** *The picture shows the holder and the microreactor clipped on the Evotip after sliding the microreactor and Evotip out of the holder for subsequent centrifugal transfer of the sample into the Evotip.*
